# Downregulation of PIK3IP1/TrIP on T cells is controlled by TCR signal strength, PKC and metalloprotease-mediated cleavage

**DOI:** 10.1101/2024.04.29.591680

**Authors:** Benjamin M. Murter, Sean C. Robinson, Hridesh Banerjee, Louis Lau, Uzodinma U. Uche, Andrea L. Szymczak-Workman, Lawrence P. Kane

## Abstract

The protein known as PI3K-interacting protein (PIK3IP1), or transmembrane inhibitor of PI3K (TrIP), is highly expressed by T cells and can modulate PI3K activity in these cells. Several studies have also revealed that TrIP is rapidly downregulated following T cell activation. However, it is unclear as to how this downregulation is controlled. Using a novel monoclonal antibody that robustly stains cell-surface TrIP, we demonstrate that TrIP is lost from the surface of activated T cells in a manner dependent on the strength of signaling through the T cell receptor (TCR) and specific downstream signaling pathways, in particular classical PKC isoforms. TrIP expression returns by 24 hours after stimulation, suggesting that it may play a role in resetting TCR signaling at later time points. We also provide evidence that ADAM family proteases are required for both constitutive and stimulation-induced downregulation of TrIP in T cells. Finally, by expressing truncated forms of TrIP in cells, we identify the region in the extracellular stalk domain of TrIP that is targeted for proteolytic cleavage.

## Introduction

Signaling pathways controlled by phosphoinositide (PI) intermediates are ubiquitous regulators of cell growth, survival and transformation (1). Key signal transducers in this pathway include various phosphatidylinositol 3-kinases (PI3K), which catalyze phosphorylation of PI species at the 3’ position of the inositol ring to produce PIP_3_ and related species. The second messenger PIP_3_ acts to recruit pleckstrin homology (PH) domain-containing proteins like Akt to the plasma membrane. In total, four different catalytic subunits (α, β, γ, δ) comprise the PI3K family, which pair with adaptor subunits that allosterically regulate PI3K activation (2–4). The major PI3K isoform linked to T cell activation is PI3Kd. The PI3K signaling pathway is a significant mediator of activation, survival and differentiation signals downstream of the TCR and CD28, integrating antigen recognition and co-stimulatory signals, respectively (5–8). Additionally, IL-2 and IL-15 cytokine receptors also activate PI3Ks to modulate T cell survival and function (9). Counter-regulating these activating signals are several phosphatases, most notably the lipid-phosphatase and tumor suppressor PTEN, which regulate these signals and tune the T cell response (10). *In vivo*, dysregulation of PI3K signaling in T cells can lead to the development of hematological malignancy or immunodeficiency (11–13).

While it has long been known that PI3Ks are activated in response to TCR for antigen and co-stimulatory molecule (e.g. CD28) engagement, the precise role of this pathway in T cell responses has been more challenging to parse. Andreatti, et al. previously demonstrated that the PI3K signaling pathway plays a more nuanced role in T cell biology (14) when compared with the critical TCR-proximal kinases (Lck, Zap70) and adaptor proteins (LAT, SLP-76). Thus, a fuller understanding of the role of this pathway in T cell biology has required *in vivo* approaches, where more subtle mechanisms can be revealed through downstream effects on T cell differentiation.

We and others recently reported that T cell activation can be modulated by a novel negative regulator of PI3K that we have referred to as <u>t</u>ransmembrane <u>i</u>nhibitor of <u>P</u>I3K (TrIP; gene name *PIK3IP1*) (15–17). Unique among known negative regulators of PI3Ks, TrIP is a transmembrane protein that contains an SH2-homology domain toward its C-terminus similar to the inter-SH2 domain of p85α/β, which are critical upstream positive regulators of PI3K catalytic subunit (p110 family) activation. Although the precise molecular mechanism by which TrIP functions is not clear, evidence suggests that TrIP specifically interacts with p85 that is recruited to phospho-Tyr residues in proteins like CD28, which in turn interferes allosterically with the activation of p110 by p85. This interaction between TrIP and the p110/p85 heterodimer has been shown to negatively regulate the catalytic activity of PI3K and inhibit downstream signaling (18).

Although initial studies on TrIP shed light on its roles in T cell activation, they were somewhat limited by the lack of a suitable monoclonal antibody (mAb) that could assay cell-surface expression of TrIP. In this study, we detail the production and validation of such antibodies, focusing on the highest sensitivity mAb clone 18E10. We have used this antibody to characterize the levels of TrIP on various immune cell types, finding that neutrophils and CD8^+^ T cells have particularly high expression. We confirm that endogenous cell-surface TrIP protein is lost upon TCR activation on CD8^+^ T cells, discovering that this downregulation is related to strength of activating TCR signal. We also are the first to report that although highly expressed in naïve cells, TrIP expression on CD8^+^ T cells does return at later time points following activation. Additionally, we have found that PKC activation downstream of TCR signaling is required to trigger the downregulation of TrIP surface expression. Furthermore, we have determined that matrix metalloproteases of the ADAM family are responsible for the downregulation of endogenous TrIP protein from activated T cells. Finally, we have discovered that this downregulation is mediated by site-specific cleavage in the stalk domain. Taken together, these studies advance our knowledge of both how TrIP is regulated at the cellular level, and how this is tied to initiation of T cell activation. Our identification of a non-cleavable form of TrIP should promote further understanding of the wider biological effects of this protein *in vivo*.

## Results

### Development of monoclonal antibodies to murine TrIP

The *in vivo* study of TrIP function has been hampered by the lack of suitable antibodies, especially against the extracellular domain of the protein. We therefore generated an Ig fusion protein by cloning the extracellular domain of TrIP into a vector encoding the Fc region from human IgG1 (TrIP-Ig; **Fig. 1A**). This construct was then used to produce recombinant TrIP-Ig by transfection into HEK293T cells. After confirming its integrity by SDS-PAGE (**Fig. 1B**), TrIP-Ig was used to immunize rats. Spleens of immunized rats were harvested and fused with myeloma cells to generate hybridomas. Supernatants from stable clones were then screened against TrIP-Ig and counter-screened against human IgG1. Supernatants from three sub-clones were then screened for the ability to recognize murine TrIP (mTrIP) by flow cytometry. Initially, HEK293T cells were transiently transfected with a Flag-tagged mTrIP (Flag-mTrIP) construct and labeled with supernatant from each anti-TrIP mAb clones. Following up with an anti-rat IgG secondary to label the clones with a fluorophore we show in **Figure 1C**, all three of these mAb’s specifically bound to cells expressing F lag-mTrIP, showing double positive staining for Flag as control. The degree of binding, however, varied by clone with 18E10 showing the most robust labeling of mTrIP based on co-labeling with an anti-Flag mAb (**Fig. 1C**). We also screened the anti-TrIP clones for cross-reactivity with human TrIP, as mouse and human TrIP are approximately 80% identical, and therefore could potentially cross-react (18). Although not seen with all three mAb’s, we found the 18E10 clone displayed significant cross-reactivity with ectopically expressed human TrIP (**Fig. 1D**). We next determined which of the two major extracellular regions, the stalk domain or the kringle domain, the novel anti-TrIP mAb’s were binding. To test this, we compared binding of the Ab’s to full-length TrIP vs. a construct engineered to lack the kringle domain, but still bearing a N-terminal Flag-tag on the stalk for detection. Despite robust Flag staining, binding of all the mAb’s was lost in the absence of the kringle domain (mTrIP-ΔKringle) (**Fig. 1E**). These data demonstrate that we have successfully generated several monoclonal anti-murine TrIP antibody clones, which bind via its kringle domain. Additionally, we found that that the most sensitive mAb (18E10) shows cross-reactivity against human TrIP.

**Figure 1.**
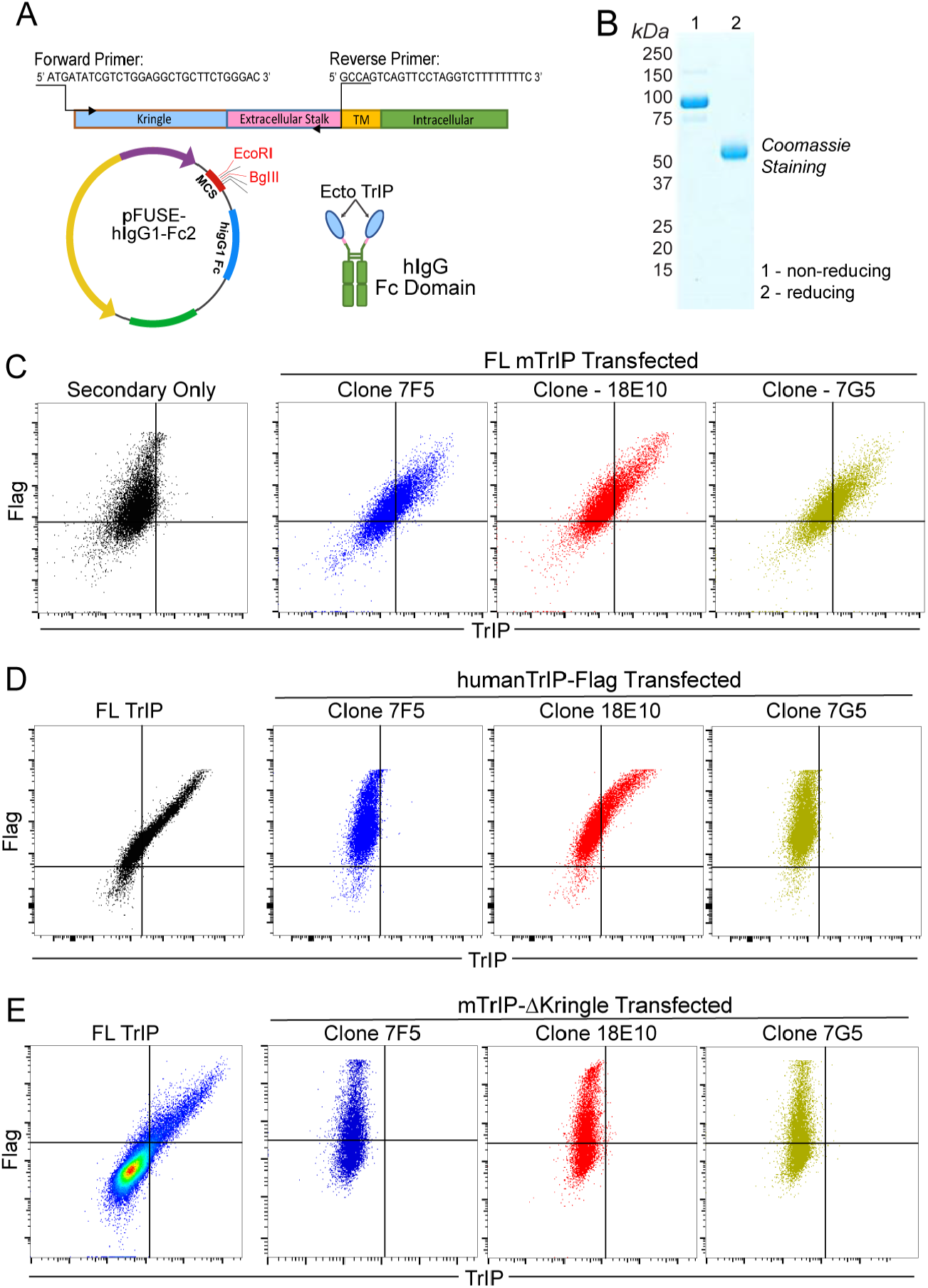
Development and validation of novel monoclonal antibodies to the ecto domain of TrIP. *A*, cloning strategy and development of the ecto-TrIP Fc-fusion protein used for immunization and antibody development. *B*, SDS-PAGE of the ectoTrIP-Ig fusion protein under reducing and non-reducing conditions. *C*, Flow cytometry data showing the binding of each anti-TrIP monoclonal antibody to a Flag-tagged WT murine TrIP construct expressed in HEK293T cells. *D*, Flow cytometry data showing staining of the clones on HEK293T cells transfected with a plasmid encoding Flag-tagged human TrIP. *E*, Flow cytometry data showing the staining of each of the anti-TrIP mAbs on HEK293T cells transfected with a Flag-tagged mTrIP construct lacking the extracellular kringle domain (ΔKringle).

### Expression of TrIP on normal murine splenocytes

We next wanted to confirm that the newly generated anti-mTrIP mAb could recognize endogenous levels of protein. Thus, we directly conjugated anti-mTrIP mAb 18E10 to Alexa Fluor 647 for use in multi-color flow cytometry. Previous reports (17, 19, 20) and public data sets (21) show that TrIP expression is not only restricted to the T cell compartment. Therefore, we incorporated the 18E10 mAb in a large spectral flow cytometry panel for murine splenocytes to assess TrIP expression more broadly. Using the ExCYT software package (22), we performed high-dimensional clustering analysis (t-SNE) of these data and were able to identify a wide range of cell types (**Fig. 2A)**. Next, we overlayed the expression pattern of TrIP as a heatmap (**Fig. 2B**), which revealed expression across cell types, with especially robust expression on CD8^+^ T cells (CD3^+^CD8^+^), whereas Treg (CD3^+^CD4^+^Foxp3^+^) and conventional CD4 (CD3^+^CD4^+^Foxp3^-^) had much lower expression in comparison (**Fig. 2C-D**). Of the major myeloid populations characterized, we did find moderate levels of TrIP expression on macrophages (CD11b^+^F4/80^+^) and the highest levels of TrIP expression were seen in the neutrophils (CD11b^+^Gr1^Hi^) (**Fig. 2C-D**). In parallel, we also labeled splenocytes from a CD8-specific TrIP knockout mouse (*TrIP*^*fl/fl*^ x E8i^Cre^), a mouse model previously generated in our lab, as a negative control (16). Thus, the 18E10 mAb robustly stained WT CD8^+^ T cells from naïve C57BL/6 mice, but not those from CD8-specific TrIP KO mice (**Fig. 2E**) confirming the staining specificity and sensitivity to endogenous levels of the protein.

**Figure 2.**
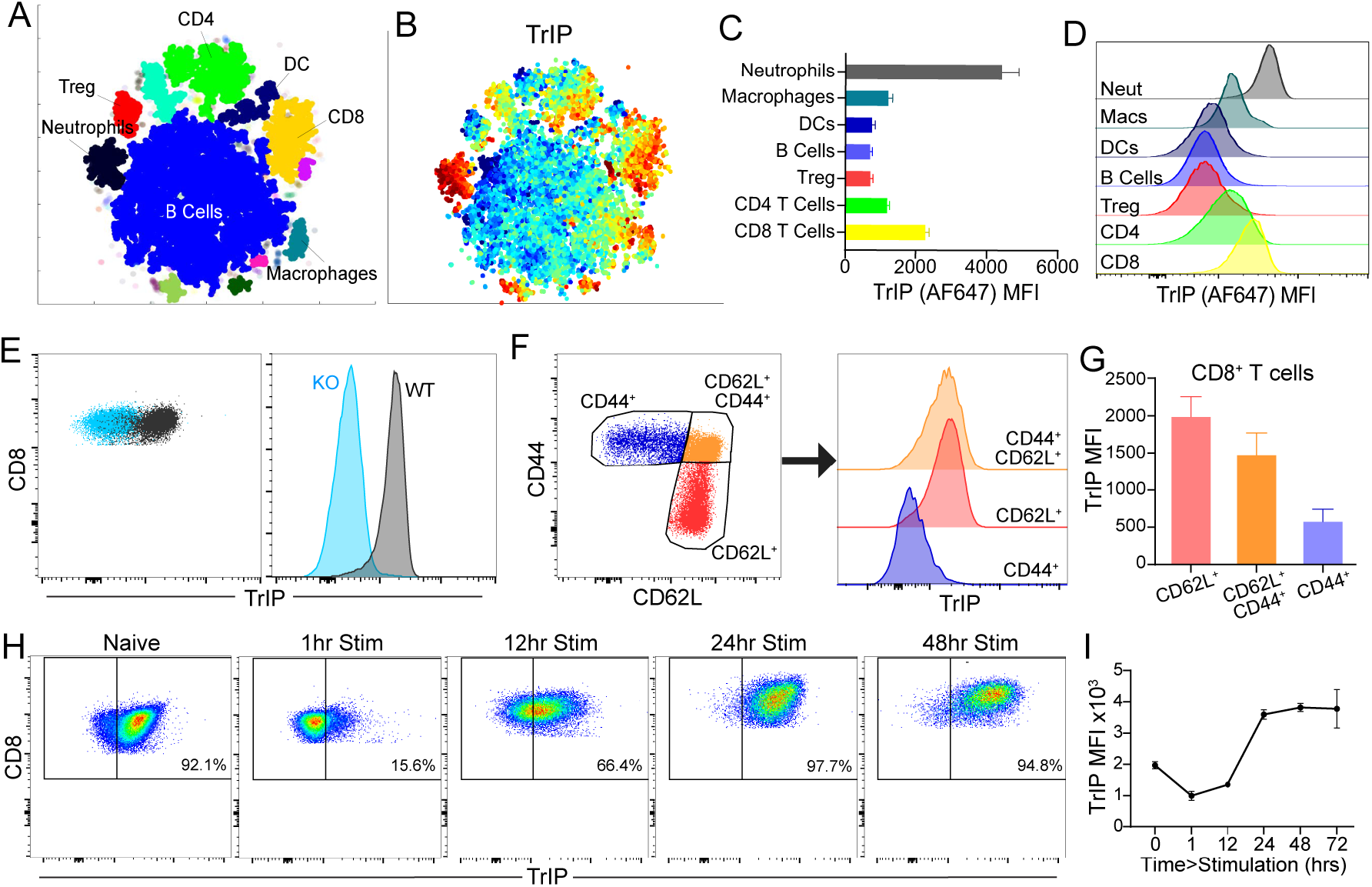
Characterization of TrIP expression on immune cells in naïve mice. *A*, ExCYT clustering analysis of flow cytometry data identifies major splenic populations including B cells (CD19^+^), CD4 (CD3^+^CD4^+^FoxP3^-^) and CD8 (CD3^+^CD8^+^) T cells, regulatory T cells (CD3^+^CD4^+^FoxP3^+^), DCs (CD11c^+^), macrophages (CD11b^+^F4/80^+^), and neutrophils (CD11b^+^Gr-1^Hi^) (22). *B*, Heatmap of TrIP protein expression overlayed across the splenic populations. *C*, Quantification of TrIP expression by MFI across each of the immune cell populations. *D*, Histograms depicting the range of TrIP expression in the indicated populations. *E*, Staining of WT vs. TrIP knockout (*Pik3ip1*^*fl/fl*^*E8i*^*cre*^) CD8^+^ T cells (CD3^+^CD8^+^) from naïve spleens as a dot plot and histogram. *F*, Gating strategy for naïve (CD62L^+^) and activated (CD44^+^) CD8^+^ T cells and corresponding histograms of their TrIP expression. *G*, Quantification of TrIP MFI in the CD8^+^ T cell subpopulations, stratified by their CD44 vs CD62L expression. *H*, Stimulation of whole WT P14 TCR Tg splenocytes with 200 ng/mL of WT gp33 peptide. TrIP was followed over 72 hrs by flow cytometry. *I*, Quantification of TrIP expression MFI on CD8^+^ T cells in the 72 hrs following activation *in vitro*.

Previous work from our lab and others has suggested that TrIP expression is high on resting cells but is downregulated upon T cell activation (15–17). In support of these data, staining of endogenous TrIP revealed that over 90% of splenic CD8^+^ T cells are TrIP^+^ (**Fig. 2E, H**). Furthermore, subsetting these cells based on their activation status revealed that TrIP was indeed expressed at much lower levels on effector (CD44^+^), vs. naïve/resting (CD62L^+^) T cells (**Fig. 2F-G**). Our previous studies using cell line expression of epitope-tagged TrIP suggested that TrIP surface expression is lost upon activation (16), so we wanted to confirm that endogenous TrIP protein behaves in a similar manner. To test this, we subjected WT P14 splenocytes to plate-bound anti-CD3/anti-CD28 stimulation, and followed the expression of TrIP with the 18E10-AF647 mAb out to 24 hours after stimulation. As shown in **Figure 2H-I**, TrIP surface expression was rapidly lost within the first two hours of stimulation and remained at very low levels for about twelve hours. To date, it has been unclear when, or even if, TrIP expression re-occurs following T cell activation. As shown in **Figure 2H-I**, we found that CD8^+^ T cells begin to re-express TrIP around 12 hours following activation and fully re-express TrIP by 24-48hrs.

### Kinetics of TrIP downregulation on activated T cells correlates with TCR signal strength

Existing data suggest that TrIP acts as a “rheostat” for initial PI3K and T cell activation downstream of the TCR. Therefore, we wanted to assess the relationship of TCR signal strength to the putative cleavage of TrIP after acti vation. Using 18E10-AF647 mAb to assay TrIP levels over time, we stimulated splenocytes from WT C57BL/6 mice with varying concentrations of plate-coated anti-CD3 plus anti-CD28 mAb’s. We found that the kinetics of TrIP loss correlated with the strength of stimulation, in a roughly semi-logarithmic fashion (**Fig. 3A-B**). Next, we tested the relative contributions of signal 1 (TCR/CD3) and sig nal 2 (CD28 co-stimulation) on the downregulation of TrIP following activation. To block CD28 co-stimulation, we pre-treated WT splenocytes with saturating amounts of CTLA4-Ig prior to stimulation with anti-CD3 mAb. These experiments suggested that the loss of TrIP does not require CD28 co-stimulation, as the degree and timing of TrIP downregulation were nearly identical with or without CD28 blockade (**Fig. 3C-D**).

**Figure 3.**
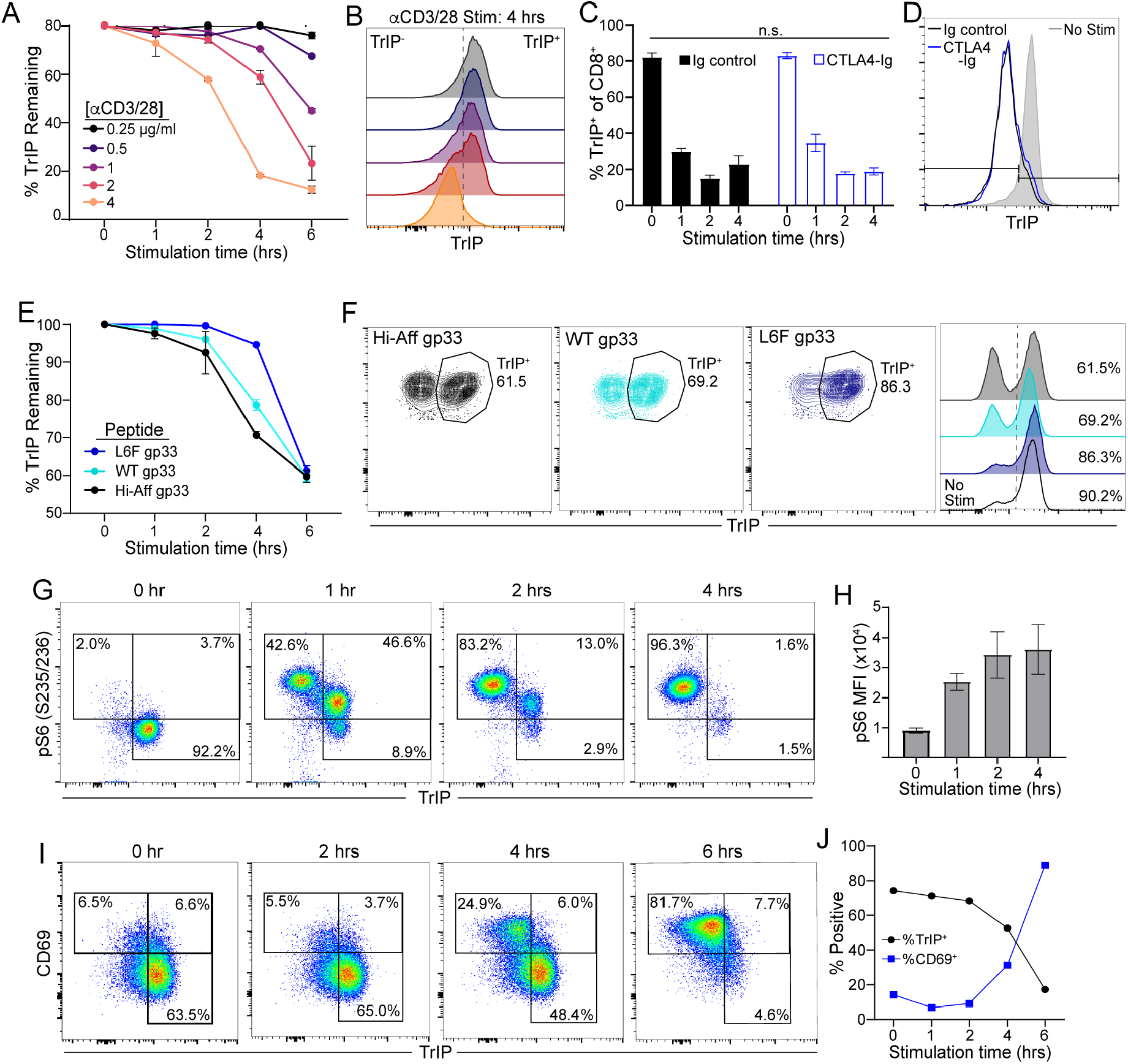
Stimulation strength-dependent downregulation of TrIP on CD8^+^ T cells. *A*, WT C57BL/6 splenocytes were stimulated with plate-coated anti-CD3 and anti-CD28 at the indicated concentrations and TrIP loss on the CD8^+^ T cells was followed over time by flow cytometry. *B*, Histogram of the TrIP expression on CD8^+^ T cells at the 4-hr timepoint of each stimulation dose. *C*, WT B6 splenocytes stimulated with 2 μg/mL of plate-coated anti-CD3 in the presence or absence of saturating amounts of CTLA4-Ig (20 μg/mL). *D*, Representative histogram overlay of CTLA4ig blockade experiment at the 1-hr timepoint. *E*, WT P14 TCR Tg splenocytes were stimulated with 100 ng/mL of the indicated cognate peptide variants (peptide affinity: Hi-Aff gp33>WT gp33>L6F gp33) and followed TrIP expression on the CD8^+^ T cells over time via flow cytometry. *F*, Representative TrIP staining of each peptide stimulation at the 4-hr timepoint, shown by both dot plot and histogram overlays. **G**, WT P14 splenocytes stimulated with 200 ng/mL of WT gp33 peptide showing staining for pS6_(S235/S236)_ vs TrIP staining in the CD8^+^ T cells following activation. *H*, Quantification of pS6_(S235/S236)_ MFI during the 4-hr course of stimulation. *I*, In the same WT P14 stimulation described in G-H, flow plots depicting CD69 vs. TrIP expression through 6 hrs of stimulation. *J*, Frequencies of total TrIP^+^ and total CD69^+^ cells. Ordinary two-way ANOVA used for statistical comparisons (p: *<0.05, **<0.01, ***<0.001, ****<0.0001).

To assess the sensitivity of TrIP loss to changes in TCR signal strength under more physiological conditions, we performed stimulation of T cells with antigen and antigen-presenting cells (APC). We thus stimulated splenocytes from P14 TCR Tg mice with cognate antigen (LCMV gp33 peptide presented by H-2D^b^) and measured levels of cell surface TrIP on CD8^+^ P14 T cells over time, by flow cytometry. As shown in **Figure 3E**, we observed rapid and dramatic loss of cell-surface TrIP upon stimulation with gp33 peptide, similar to what we had observed after stimulation with anti-CD3 mAb. Importantly, this system also allowed us to test the effects of altered gp33 peptides with different affinities for the P14 TCR (23). Shown in **Figure 3E-F** are the results with two such peptides, with the higher affinity peptide (HiAff gp33) triggering the greatest degree of TrIP downregulation (>30%) by four hours after stimulation. By contrast, a lower affinity peptide (L6F gp33) caused very little TrIP loss after four hours (<5%). Thus, these data demonstrate that peptide/MHC affinity for the TCR is tightly linked with the kinetics of TrIP downregulation.

Although the functional relevance of acute cell-surface TrIP downregulation is not completely clear, such down-modulation of a negative regulator like TrIP may help to constrain early T cell activation, as suggested in previous publications from our group and others (16, 17). Consistent with this notion, we found that WT P14 CD8 T cells activated with cognate peptide seem to progress through three distinct stages of early T cell activation, distinguished by their relative expression of TrIP and phosphorylation of the S6 ribosomal subunit (pS6) (**Fig. 3G**). Starting with a naïve (i.e. TrIP^+^) population of CD8^+^ T cells, we found that after one hour of stimulation with 200 ng/mL of WT gp33 peptide, a small proportion (<10%) of the cells had yet to undergo activation, indicated by a lack of pS6 staining. Somewhat unexpectedly, we found a distinct population of pS6-intermediate (pS6^int^) cells which still maintained their TrIP expression. This intermediate pS6 intensity is consistent with the regulatory activity that TrIP plays during early T cell activation, as loss of TrIP coincides with a secondary increase in pS6 signal intensity that then persists beyond four hours of stimulation (**Fig. 3H**). These data further suggest that the loss of TrIP is not a pre-requisite for active PI3K signaling, but rather may act to modulate PI3K signal intensity at the early stages of CD8^+^ T cell activation. To provide additional functional context to TrIP expression following T cell activation, we assessed whether there was a temporal association between TrIP downregulation and expression of a well-established early indicator of T cell activation, CD69. As shown in **Figure 3I**, we did in fact observe a tightly associated inverse correlation between loss of cell-surface TrIP and expression of CD69 (**Fig. 3J**).

### Role of TCR-dependent signaling pathways in acute downregulation of TrIP

Next, we sought to understand which TCR signaling pathways regulate the stimulation-induced turnover of TrIP surface expression. Initially, we hypothesized that the pS6 upregulation that occurs while still maintaining TrIP expression that we saw previously (**Fig. 3G**) may be a prerequisite for its loss. This idea suggests that PI3K signaling itself may act in a feed-forward loop, directly leading to TrIP downregulation. To investigate this, we took a pharmacological approach, using well-defined inhibitors to target various PI3K pathway members, including PI3K itself (IC87114), Akt (MK2206), and mTORC1 (rapamycin). Using WT P14/cognate gp33 peptide stimulation, we pre-treated with the indicated inhibitors for 30 minutes prior to the addition of a 200ng/ml dose of WT gp33 peptide. We found that treatment with each of these inhibitors resulted in a modest (∼10%) higher expression of TrIP up to two hours after stimulation, compared to the control (**Fig. 4A-B**). While these inhibitors did modestly slow the loss of TrIP, they did not seem to impact baseline expression levels, despite drug exposure for the entire assay (“0hr” samples) nor do they lead to persistence at later timepoints (4hr timepoints). As expected, inhibition of this pathway drastically reduced pS6_(235/2 36)_ levels, compared to the vehicle controls at the two-hour timepoint (**Fig. 4C-D**). Despite the reduction in pS6 caused by each of the drugs, it is notable that although T cells treated with MK2206 or rapamycin displayed lower pS6_(235/236)_ levels, that did not translate to increased TrIP expression, in contrast to what we seen in cells treated with IC87114 (**Fig. 4C-D**). Taken together, these data do not support the hypothesis that TrIP surface expression is regulated via an autocrine PI3K-pathway related mechanism.

We next expanded the scope of these experiments to include inhibitors targeting a broader ran ge of signaling pathways. This included inhibitors of calcium signaling (FK-506; calcineurin inhibitor), ERK (U0126; MEK1/2 inhibitor), and dasatinib (Src family kinase inhibitor). Perhaps unsurprisingly, the potent inhibition of upstream kinases by dasatinib (which would include Lck) resulted in broad inhibition of TCR signaling and significant preservatio n of TrIP expression (>50%) at two hours after stimulation (**Fig. 4E-F**). Although, importantly, dasatinib treatment at this dose did not completely prevent T cell activation and TrIP loss, as we still observed complete TrIP loss at the four-hour timepoint, similar to the vehicle group (**Fig. 4E-F**). Moving further downstream of the TCR, we found that inhibition of either calcineurin or MEK1/2 activity under these conditions also led to signific ant preservation of TrIP expression (at least 60% and 40% of starting levels, respectively) in CD8^+^ T cells following two hours of stimulation, compared to around 18% in the untreated control (**Fig. 4E-F**). While these compounds had statistically significant effects on TrIP expression, they did not detectably impact pS6_(235/236)_ levels (**Fig.4G,H**). These results suggest that while calcium and ERK-MAPK signaling play critical roles in regulating TrIP downregulation, there is also a contribution, although more minor, of the PI3K pathway itself.

**Figure 4.**
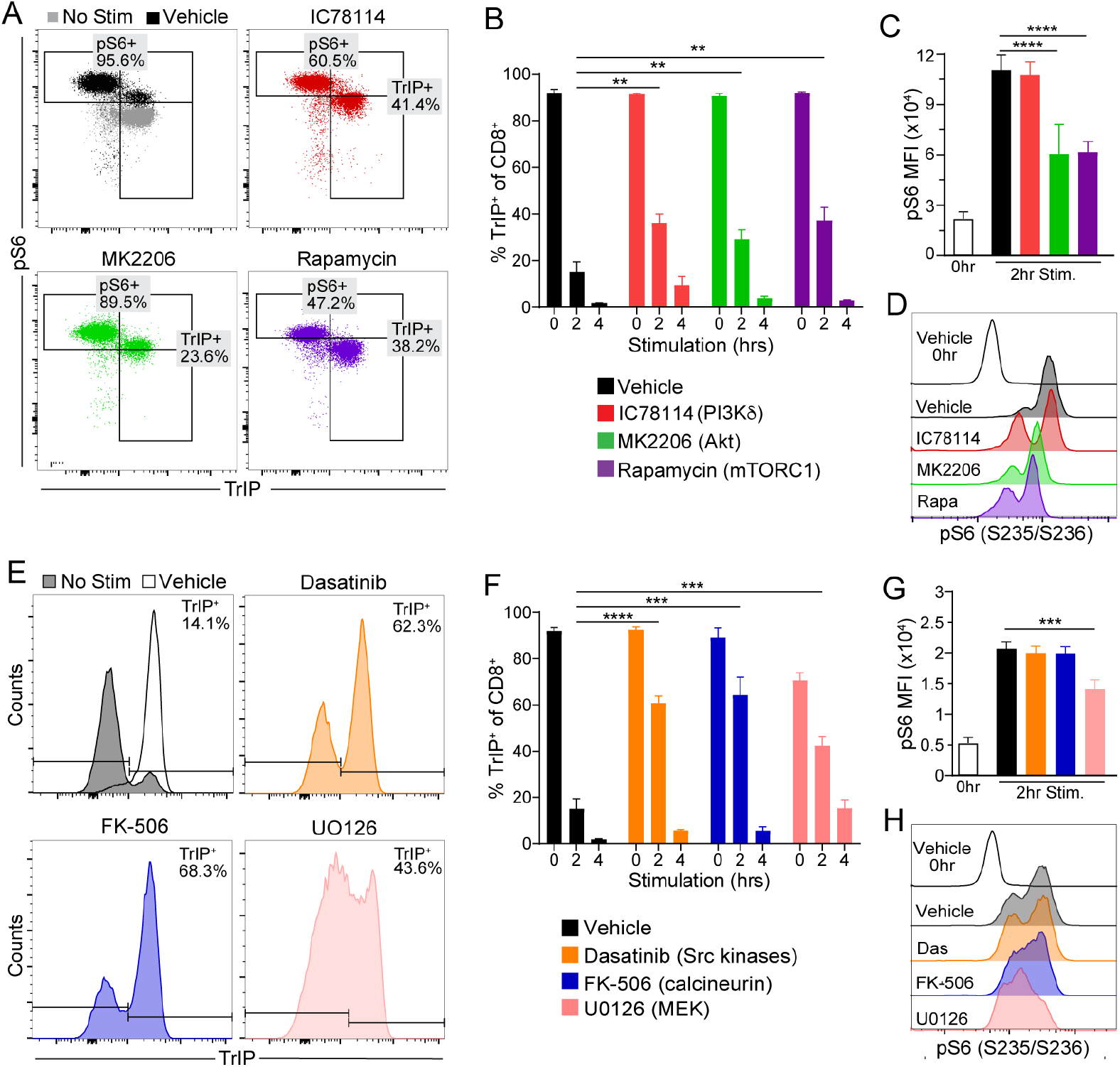
The PI3K pathway is dispensable for TrIP downregulation. WT P14 splenocytes were pre-treated for 30 minutes with the indicated inhibitors and then stimulated with 200 ng/mL cognate WT gp33 peptide, stil in the presence of inhibitor. TrIP loss on the CD8^+^ T cells was followed over time by flow cytometry. *A*, Dot plots showing pS6_(S235/S236)_ vs. TrIP expression on the CD8^+^ T cells at the 2-hr timepoint. *B*, Quantification of TrIP expression at each timepoint for each treatment *C*, Quantification of pS6_(S235/S236)_ MFI at the 2-hr timepoint, with 0hr values as comparison. *D*, Representative histograms of pS6_(S235/S236)_ staining at the 2-hr timepoint, with 0-hr histogram as comparison. *E*, Representative histograms of TrIP expression at the 2-hr timepoint following stimulation. *F*, Quantified summary of TrIP expression at each timepoint for each treatment. *G*, Quantification of pS6 levels at the 2-hr timepoint, with 0-hr values as comparison. *H*, Representative histograms of pS6 staining at the 2-hr timepoint with 0-hr histogram as comparison. Ordinary two-way ANOVA was applied for statistical comparisons (p: *<0.05, **<0.01, ***<0.001, ****<0.0001).

In addition to calcium (Ca^2+^) and PIP_3_, another critical second messenger downstream of TCR signaling is diacylglycerol (DAG), which functions in part to activate several isoforms of protein kinase C (PKC). The PKC family of serine/threonine kinases comprises three major classes based on their physiological activators. The classical PKCs (α, βI, βII, and γ) and the novel PKCs (δ, ε, η, and θ) both require diacylglycerol (DAG) for activation, but while the classical PKCs also require Ca^2+^ the novel PKCs do not (24). By contrast, so-called atypical PKCs (ζ and λ/ι) require neither DAG nor Ca^2+^ for activation (24). Since PKCs are connected to both calcium and ERK signaling, we hypothesized that one or more PKC family members may play a key role in regulating TrIP downregulation.

Given the complexity of the PKC family, we started our survey with pan-PKC inhibitors, including bisindolylmaleimide I (Bis I), sotrastaurin, and Gö6983. Thus, we found that broad pharmacological inhibition of PKC activity resulted in nearly complete prevention of TrIP downregulation from P14 T cells at two hours after peptide stimulation (**Fig. 5A-B**). Although these inhibitors share similar activities against the various PKC family members, they resulted in somewhat different kinetic effects on downregulation of TrIP and induction of pS6 (**Fig. 5C-D**), which could be due to differential off-target effects of the compounds. To better discriminate between the role of classical (α, βI, βII, and γ) and novel PKCs (δ, ε, η, and θ) in TrIP downregulation we treated T cells with Go6976, a PKC inhibitor with high selectivity for classical PKC’s and little activity against PKCθ (25), the predominant PKC activated by the TCR (26, 27). Treatment with Go6976, similar to the other PKC inhibitors, also led to significant inhibition of TrIP downregulation, compared to vehicle control (**Fig. 5A-B)**). These data suggest that PKCθ activity alone is insufficient to trigger comparable degrees of TrIP downregulation, compared to controls without inhibitor. Together, these data suggest that PKCs play a major role in regulating the surface downregulation of TrIP following TCR stimulation. While both classical and novel PKCs may contribute to TrIP cleavage, our data suggest that classical PKCs play a more predominant role in this process.

**Figure 5.**
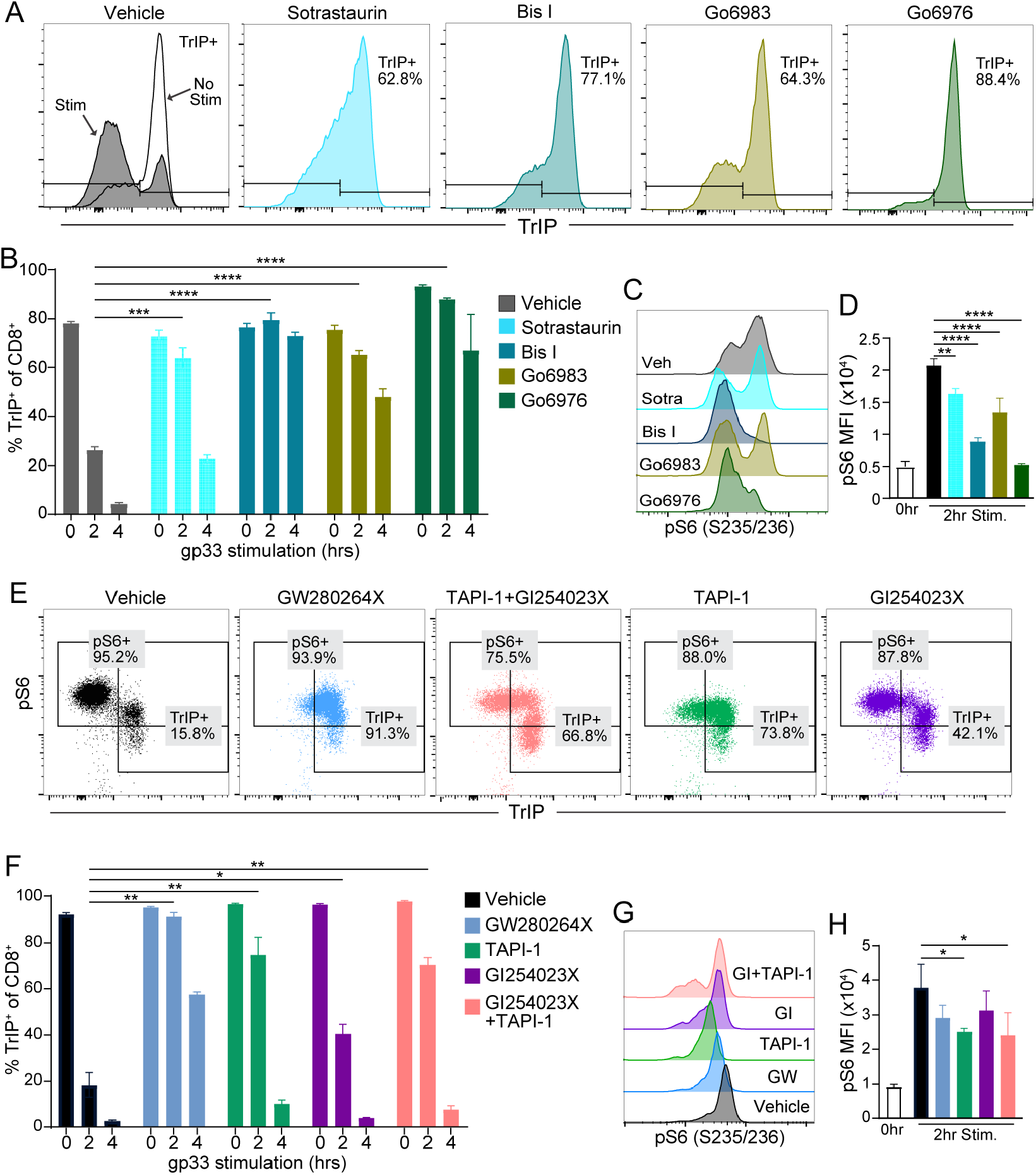
PKC and ADAM17 activity are required for downregulation of cell-surface TrIP. WT P14 splenocytes were stimulated with 200 ng/mL WT gp33 peptide in the presence of the indicated inhibitors. TrIP loss on the CD8^+^ T cells was followed over time by flow cytometry. *A*, Flow plots of TrIP staining in each PKC inhibitor treatment at 2-hr timepoint following stimulation. *B*, Quantification of TrIP expression at each timepoint for each PKC inhibitor treatment. *C*, Representative histograms of pS6_(S235/S236)_ staining at the 2-hr timepoint. *D*, Quantification of pS6_(S235/S236)_ MFI at the 2-hr timepoint. *E*, Representative flow plots pS6_(S235/S236)_ vs. TrIP expression treatment at the 2-hr timepoint in the presence of ADAM inhibitors. *F*, Quantification of TrIP expression across each ADAM inhibitor treatment. *G*, Representative histograms of pS6_(S235/S236)_ staining at the 2-hr timepoint. *H*, Quantification of pS6(S235/S236) MFI at the 2-hr timepoint. Ordinary two-way ANOVA statistical analysis applied for statistical comparison (p: *<0.05, **<0.01, ***<0.001, ****<0.0001).

### Role of metalloproteases in acute TrIP downregulation

While the above data further define the cellular signaling pre-requisites for TrIP downregulation, we have yet to fully understand the direct mechanism by which TrIP is lost. Shedding of surface proteins by proteolytic cleavage has been shown to play numerous immunological roles, with the ADAM (A disintegrin and metalloproteinase domain-containing) family of proteases playing prominent roles (28–30). In fact, our initial studies using expression of Flag-tagged murine TrIP constructs in T cells (16) revealed that the inhibitor GW280264X (“GW,” a dual inhibitor of ADAM10/ADAM17) prevented stimulation-induced downregulation of TrIP (16). Thus, we sought to evaluate if endogenous murine TrIP is regulated in a similar manner. Expanding on our previous work, we set out to determine the relative contributions of ADAM10 vs ADAM17 in the cleavage of TrIP from the surface. In addition to the ADAM10/17 dual inhibitor (GW), we also utilized GI254023X (GI) an ADAM10 inhibitor with 100-fold selectivity for ADAM10 over ADAM17. There are limited specific inhibitors for ADAM17 that do not interfere with ADAM10, but using TAPI-1 we could at least preferentially target ADAM-17, as it has a 10-fold lower potency for ADAM10. As shown in **Figure 5E-F**, potent inhibition of both ADAM10/17 (using GW) resulted in significant preservation of TrIP expression following stimulation. ADAM17 inhibition by TAPI-1 appeared to more significantly preserve cell-surface TrIP (20% loss) compared to the ADAM10-inhibited (GI) group (40% loss) (**Fig. 5E-F**). Furthermore, when we combined these compounds, the effect was not additive, instead mimicking inhibition with TAPI-1 alone (**Fig. 5E-F**). Finally, these experiments reinforced our understanding of negative regulation of PI3K by TrIP, as conditions in which TrIP expression was maintained also had lower levels of pS6 induction (**Fig. 5G-H**), consistent with maintenance of TrIP-mediated PI3K inhibition. Together, these data confirm that downregulation of cell-surface TrIP expression in CD8^+^ T cells is mediated, at least in part, by ADAM10 and ADAM17.

Given the limitations of the pharmacological tools, we next sought to deconvolute the relative contributions of ADAM10 vs. ADAM17 to the cleavage of TrIP from the surface, as well as further define the role of PKC activity in this pathway. We thus used CRISPR/Cas9 to target ADAM10, ADAM17, and PKCθ. While both pharmacologic and genetic approaches can result in off-target effects, additional genetic confirmation of the above data would strengthen our understanding of the mechanisms of TrIP expression regulation. Two crRNAs per gene were designed to target to the genes encoding PKCθ, ADAM10 and ADAM17 (see Methods). There are several challenges with targeting genes via CRISPR in naïve T cells, as most protocols involve prior or concurrent activation of the T cells. We felt it was critical to use naïve CD8^+^ T cells for these experiments, to be consistent with the experiments described above and given that stimulation through the TCR causes acute downregulation of TrIP. Based on a published protocol, we used beads to isolate naïve CD8 T cells and incubated them in rIL-7 for 24 hours prior to nucleofection (31, 32). These T cells were then cultured in IL-7 for an additional five-to-seven days, prior to stimulation, to allow for turnover of message and protein.

Using this approach, we achieved robust down-regulation of ADAM10 by day five in culture, as shown by the flow plots in **Figure 6A**. We discovered CRISPR knockout of ADAM10 in naïve CD8 T cells resulted in a significant increase in baseline TrIP expression by mean fluorescence intensity (**Fig. 6A-B**), suggesting that ADAM10 plays a role in constitutive shedding of endogenous TrIP. Additionally, ADAM10 knockout led to a modest, but statistically significant, attenuation of the overall degree of TrIP downregulation following TCR stimulation, at the level of overall TrIP MFI (**Fig. 6C-D**). We observed a similar effect when quantifying the percentage of cells falling into the TrIP^+^ gate after stimulation (**Fig. 6I**). These data provide additional evidence that ADAM10 plays a role in both constitutive and inducible TrIP downregulation, via cleavage. These data are in line with the literature, as ADAM10 has been shown to be responsible for both constitutive and also induced and/or regulated shedding of other proteins as well (33, 34). Next, we sought to knock out ADAM17 in naïve T cells using the same approach. However, despite designing multiple guide RNA’s and other protocol modifications, we were not able to detectably alter the expression of ADAM17 in naïve CD8^+^ T cells following CRISPR-targeted knockout (not shown). While this could be due to multiple reasons, we believe ADAM17 protein turnover and/or degradation had not occurred, despite testing for ADAM17 knockout up to 10 days following electroporation. This idea is supported by the literature, as lysosomal degradation of mature ADAM17 has been shown to require PKC activation (35), consistent with the lack of ADAM17 turnover that we observed.

**Figure 6:**
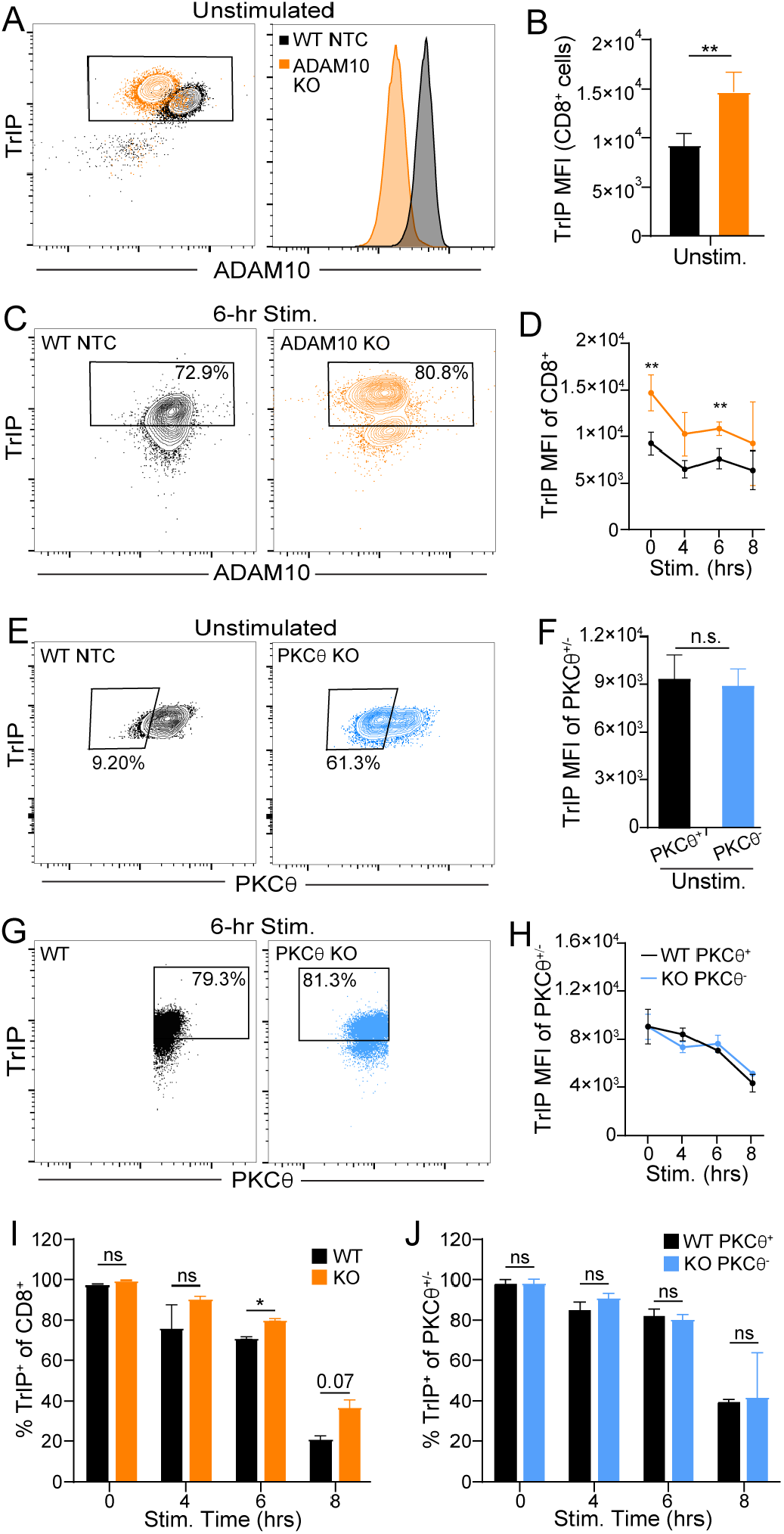
CRISPR-mediated knockout in naïve CD8 T cells confirms a role for ADAM10 in TrIP downregulation and suggests that PKCθ is dispensable. Naïve T cells were transfected with Cas9 RNP’s targeting ADAM10 or PKCθ, then maintained in rIL-7 for 7 days *in vitro*, followed by stimulation with α-CD3 mAb. *A-B*, Knockout efficiency of ADAM10, shown by flow cytometry (A) and histogram (B), which indicate an increase in basal TrIP expression before stimulation. TrIP expression was assessed on day 7 after nucleofection, prior to stimulation. *C-D*, Expression of TrIP and ADAM10 after stimulation of the indicated cells, at the indicated time points. Samples were gated on total CD8^+^ cells. *E-F*, Expression of PKCθ and TrIP at day 7 after nucleofection, and before stimulation, shown by flow cytometry (E) and histogram (F). MFI data in panel F are from the PKCθ^+^ or PKCθ^−^ gate in panel G for WT and KO cells, respectively. *G-H*, Expression of TrIP and PKCθ after stimulation of the indicated cells, at the indicated time points. MFI data in panel H are from the PKCθ^+^ or PKCθ^−^ gate in panel G for WT and KO cells, respectively. *I-J*, The percentage of TrIP^+^ cells at the indicated time points after stimulation. Cells in panel I (ADAM10 KO) were gated on total CD8^+^ cells; cells in panel J were gated on PKCθ^+^ or PKCθ^−^ cells in WT and PKCθ KO conditions, respectively. Ordinary two-way ANOVA was used for statistical comparisons (p: *<0.05, **<0.01, ***<0.001, ****<0.0001).

We next turned our attention back to PKCθ, using the CRISPR approach. Thus, we were able to achieve a marked, although incomplete (∼60%), reduction in PKCθ staining (**Fig. 6F**). However, we did not observe an increase in the baseline level of TrIP upon PKCθ KO (**Fig. 6G**) like we saw in the ADAM10 KO, even when gating specifically on PKCθ^+^ vs. PKCθ^-^ cells, suggesting that PKCθ is not required for the constitutive shedding of TrIP. Consistent with this finding, we also did not obseve any effect of PKCθ knockout on stimulation-dependent TrIP downregulation (**Fig. 6G-H, J**), again even when gating on PKCθ^+^ vs. PKCθ^-^ cells in the WT and KO samples, respectively. These data indicate that, although PKCθ is the major isoform of PKC linked to many TCR signaling-dependent events, other PKC family members play more dominant roles in the downregulation of TrIP, consistent with the above pharmacological data.

### Structural requirements for TrIP downregulation

Next, we wanted to understand where in the ecto domain of TrIP the ADAM-mediated cleavage occurs. We focused on the 76-amino acid long connecting ‘stalk domain’ linking the extracellular kringle domain to the transmembrane and intracellular domains (**Fig. 7A**). Assessment of the putative structure of this protein in AlphaFold (36, 37) suggests that the extracellular ‘stalk’ domain region of TrIP is unstructured in nature (**Fig. 7A**). Initially, we explored *in silico* prediction methods (e.g., iProt-Sub) to identify predicted metalloprotease cleavage sites (38). This analysis revealed multiple possible sites of cleavage, which spanned a wide range of the stalk domain (**Fig. 7A**), and thus did little to narrow down the region(s) where cleavage might occur.

**Figure 7.**
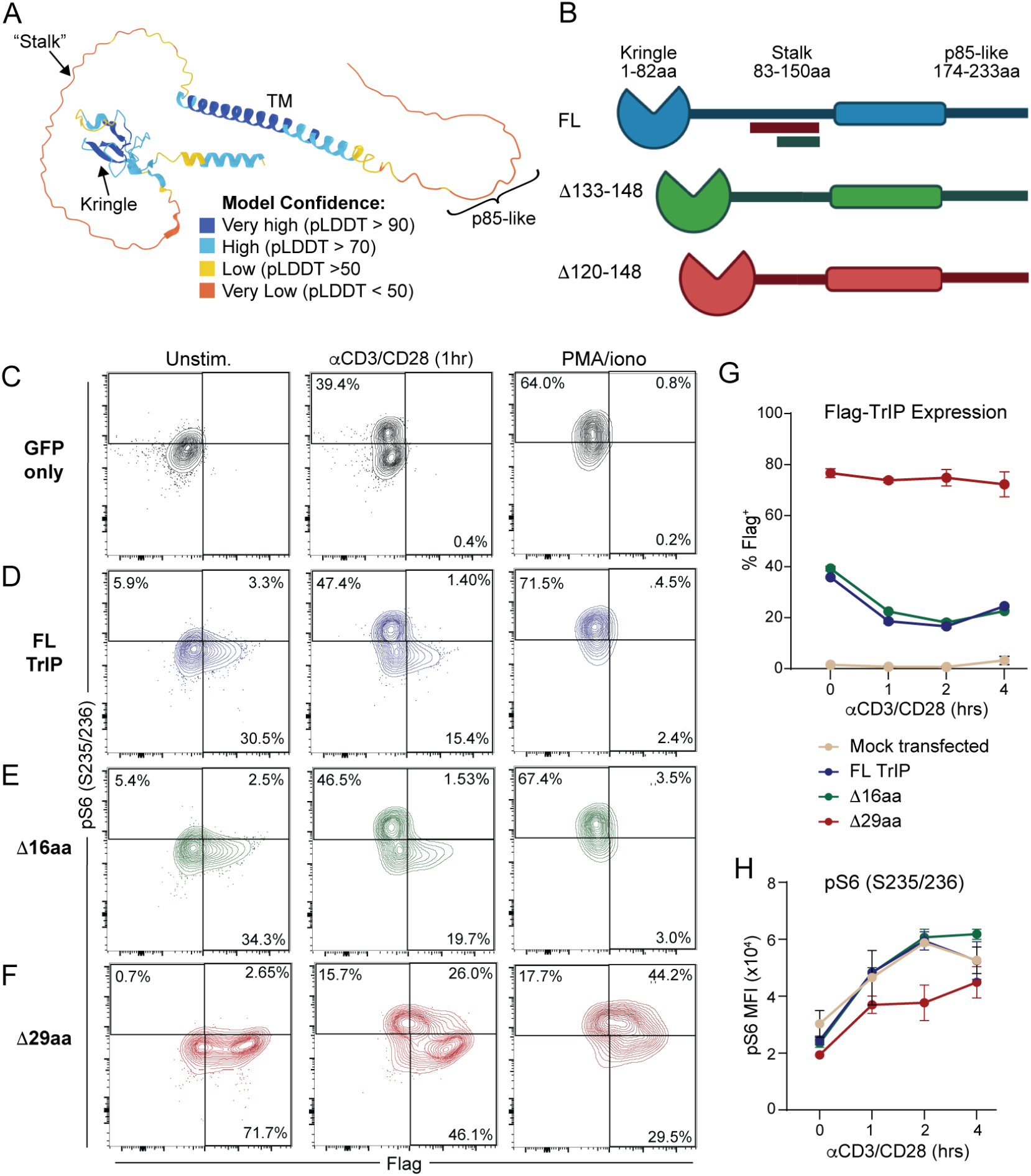
Truncation of the extracellular stalk renders TrIP resistant to cleavage and dampens PI3K signaling. D10 T cells were co-transfected with pMaxGFP alone or together with the indicated TrIP plasmid constructs. Twenty four hours following transfection, cells were stimulated with 10 μg/mL plate-coated anti-CD3 + anti-CD28 or PMA/ionomycin (50 ng/mL and 500 ng/mL, respectively). *A*, AlphaFold sourced predicted crystal structure of TrIP protein, indicated is the model confidence by color. *B*, Diagram depicting the two truncation mutants used in these studies. All flow plots show cells gated on GFP^+^. *C*, Control GFP-only transfected D10 cells (GFP only) depicted without stim, following 1-hr of αCD3/CD28 stimulation, and following 1-hr of PMA/ionomycin stimulation. *D*, Co-GFP+WT TrIP (FL TrIP) transfected D10 T cells depicted without stimulation, following 1-hr of αCD3/CD28 stimulation, and following 1-hr of PMA/ionomycin stimulation. *E*, D10 T cells co-transfected with GFP and the Δ16aa mutant of TrIP (16aa del’n) depicted without stim, following 1-hr of αCD3/CD28 stimulation, and following 1-hr of PMA/ionomycin stimulation. *F*, D10 T cells co-transfected with GFP + the Δ29aa mutant of TrIP (29aa del’n) depicted without stimulation or following 1-hr of stimulation with αCD3/CD28 or PMA/ionomycin. *G*, Quantification of TrIP-Flag (%Flag^+^) in transfected D10 cells following αCD3/CD28 stimulation. *H*, Quantification of pS6_(235/236)_ MFI in the transfected D10 cells following αCD3/CD28 stimulation.

The shortcomings of these types of prediction are likely due to the somewhat promiscuous target specificity of ADAM metalloproteases (39, 40). Previous reports suggest that ADAM mediated cleavage typically occurs in sites proximal to the cell membrane (41), so we focused on that area of TrIP. We therefore made a series of internal deletion mutants to confirm that both; cleavage is the mechanism by which cell-surface TrIP is downregulated, and to create a non-cleavable TrIP construct for future functional studies. The sites of these deletions are indicated in **Figure 7B**; each construct also included a Flag-tag for detection. Thus, we found that when we stimulated D10 T cells expressing both WT and our truncated TrIP constructs, deletion of just a sixteen amino acid sequence (Δ16aa) was not sufficient to prevent cleavage/loss (**Fig 7D-E**). By contrast, there was no detectable cleavage of a TrIP construct with a more substantial 29 amino acid deletion (Δ29aa; **Fig. 7F**). Corroborating earlier results, the inability to cleave Δ29aa TrIP construct from the cell surface led to increased basal TrIP expression (**Fig. 7G**), similar to the ADAM10 knockout data discussed above. It is still unclear whether this is due to enhanced production or processing of the transfected protein or whether it might be the result of diminished steady-state cleavage of TrIP.

As described above, our data suggested that PKC activation is required for downregulation of cell-surface TrIP, so we also stimulated transfected D10 T cells with phorbol 12-myristate 13-acetate (PMA), which directly activates multiple PKC isoforms (35, 42). The potent stimulation mediated by PMA led to near complete loss of both WT and Δ16 TrIP within one hour of treatment. Strikingly however, the Δ29aa mutant was still resistant to downregulation, even in the presence of PMA (**Fig. 7F-G**). Consistent with our previous study (16) and data shown above, expression of the cleavage-resistant Δ29 form of TrIP led to a decrease in pS6 staining in TC R-stimulated D10 T cells (**Fig. 7F**). Together, these data expand our understanding of TrIP regulation, showing that TrIP cleavage from the surface by ADAM17 (TACE) occurs in the stalk region, between residues 120 and 133, and that in the absence of TrIP downregulation, partial suppression of the PI3K/Akt pathway is maintained.

## Discussion

In this study, we expand on the understanding of TrIP, a transmembrane protein comprised of an extracellular protein-protein interaction kringle domain (43, 44), and an intracellular ‘p85-like’ domain (15, 16, 18). The p85-like domain of TrIP has been shown to interact with the p85/p110 heterodimer of PI3K and negatively regulate its catalytic activity, although this is not a complete inhibition (18). Furthermore, expression of TrIP in T cells has been shown to negatively regulate their activation and that its deletion can enhance their inflammatory activities (15, 16). While gaining valuable insight into fundamental TrIP biology, these previous studies were hindered by the lack of availability of a robust anti-murine TrIP monoclonal antibody and thus relied on cell line expression systems and knockouts to characterize its function. In this study, we describe the production and validation of three novel anti-mTrIP IgG monoclonal antibodies capable of recognizing endogenous murine TrIP protein for detection by flow cytometry. We also noted the high sequence homology (>80%) between mouse and human TrIP, and while not all mAb clones were capable of cross-reactivity we do find that the top clone (18E10) can detect both human and murine TrIP. Each of the three clones appears to stain TrIP via its kringle domain, since deletion of that domain completely abrogated their staining.

Consistent with publicly accessible gene expression data, splenic T and B lymphocytes demonstrated some of the highest levels of TrIP protein expression. Nonetheless, we also observed relatively high expression of TrIP on neutrophils, something that has not yet been reported in the literature. Furthermore, upon subsetting the CD8 T cells based on their CD44 vs. CD62L expression, we found that the naïve CD62L^+^ cells comprise the majority of the TrIP-expressing cells, whereas it was not detectable on CD44^+^ cells. We further showed that this endogenous TrIP expression on naïve CD8^+^ T cells is lost within the first few hours after TCR stimulation, confirming our previous studies with the D10 helper T cell line (16). Building upon these data, we found that TrIP expression is lost following stimulation with anti-CD3 mAb or peptide antigen in a dose-dependent manner. Furthermore, this loss appears to be largely independent of the “signal 2” (CD28) contribution, as blockade with CTLA4-Ig had little-to-no effect. While having confirmed previous studies that suggested TrIP expression would be found on naïve T cell populations and lost upon activation, it was unclear when, or even if, TrIP may be re-expressed following T cell activation. Our results show that although TrIP expression does remain low for up to twelve hours following activation, T cells do begin to re-express TrIP by 24 hours, at least *in vitro*. This re-expression of TrIP suggests it may play a previously unexplored role in longer lived populations like memory T cells, a subject that is worthy of further study.

The ability to track endogenous TrIP protein on T cells has given us a new understanding of how TrIP expression may coordinate early T cell activation signals. Ours is the first study, that we know of, to describe an early activation state in CD8^+^ T cells during which TrIP is still expressed and there is an intermediate level of PI3K activation (as read-out by pS6). This temporally correlated series of signaling events adds to our understanding of the complex role of TrIP in early T cell activation. Along with the kinetics of S6 phosphorylation, we also found that upregulation of CD69, a well-characterized marker of early T cell activation (45), was inversely correlated with the downregulation of TrIP.

We also conducted a thorough study of the relationship of TrIP downregulation to TCR signaling. While TCR ligation triggers multiple signaling pathways, we initially focused on the PI3K pathway, as we had observed a relationship between the loss of TrIP and the upregulation of pS6. By pharmacologically targeting various members of the PI3K pathway, we found that regardless of which stage of the pathway was inhibited, each resulted in small but consistent effects, i.e. slower kinetics of TrIP downregulation. These relatively minor effects may be a result of lower available PIP_3_ and/or active Akt, which can participate in parallel signaling pathways (46). For example, PIP_3_ promotes the activation of Itk, leading to enhanced PLC-γ1 activation and Ca^2+^ signaling; in addition, calmodulin is known to compete with Akt for PIP_3_ (46). Thus, this led to our expansion of the investigation to other pathways and we found that inhibition of enzymes more proximal to calcium signaling (calcineurin) or ERK activity (MEK1/2) had much greater impacts on the kinetics of TrIP loss.

The above data also led us further explore the roles of the protein kinase C (PKC) family of enzymes, as some members of this family are activated in a Ca^2+^-dependent manner and are known to promote activation of MAPK signaling (29, 47). We found that broad inhibition of PKC activity significantly inhibits the downregulation of TrIP following T cell activation, suggesting a major role in directing its cleavage from the cell surface. Although the novel isoform PKCθ is thought to be the major isoform linking TCR signaling to downstream MAPK and NF-κB signaling (48, 49), TCR stimulation is associated with activation of classical and other PKC isoforms (50). Employing Go6976, an inhibitor selective for classical PKC’s (thus leaving PKCθ activity largely intact), we again observed robust inhibition of TrIP loss following stimulation. These data suggest a primary role for classical PKCs in triggering TrIP cleavage, with the caveat that there are additional off-target effects of this compound (e.g., on other calcium-dependent kinases) (25). To more directly address the role of PKCθ, we used CRISPR-mediated knockout in naïve CD8^+^ T cells. While we were able achieve a significant degree of knockout of PKCθ in the majority (>60%) of naïve T cells, there was no significant effect on TrIP expression before or after stimulation, even when gating specifically on cells lacking PKCθ expression. In aggregate, our pharmacological and CRISPR knockout data suggest that the classical PKC isoforms are primarily responsible for inducible downregulation of TrIP in T cells.

Building on previous studies, obtained phamacological evidence for ADAM10 and ADAM17 being the major proteases responsible for downregulation (likely via cleavage) of endogenous TrIP in CD8^+^ T cells. While we were able to corroborate the role of ADAM10 via CRISPR-mediated knockout, the contribution of ADAM17 was difficult to define with this approach. Nonetheless, together our findings are consistent with reports showing extensive overlap between ADAM10 and ADAM17 substrates (51–53). The cleavage of cell surface proteins by ADAM proteases typically occurs in close proximity to the cells plasma membrane. Thus, to better define the site of TrIP cleavage, we produced TrIP constructs with truncated “stalks.” Dramatically, removal of a 29-amino acid region of the extracellular domain (residues 120-148) completely abrogated the downregulation of TrIP following activation, while a shorter deletion of this same region had no detectable effect. These data suggest that the cleavage site lies within that region, although it is formally possible that shortening the length of this region results in steric hindrance of activated ADAM access to the cleavage site. Of note, the non-cleavable (29AA deletion) construct displayed higher baseline expression of TrIP when compared to WT constructs, suggesting that TrIP is subject to low rates of basal shedding, even in the absence of TCR stimulation. We also observed an increase in basal TrIP expression in ADAM10 KO T cells, suggesting that constitutive shedding of TrIP may be driven (at least in part) by ADAM10 cleavage in this region. Further investigation is needed to fully define the specific cleavage site and to understand the role that shedding of TrIP has on *in vivo* T cell activation and differentiation, similar to studies defining the role of other immunomodulatory molecules such as Lag3 and Tim-3 (52, 54).

We have yet to directly address the fate of the TrIP cytoplasmic tail following ecto domain cleavage. Thus, the cytoplasmic portion of TrIP contains a “p85-like domain,” which is reported to inhibit the catalytic activity of PI3K (16). Of note, none of the commercially available antibodies against the C-terminal portion of TrIP were suitable for these studies, as they did not yield specific or robust enough signals when tested. However, our previous work employed a series of Flag-tagged TrIP constructs in cell line expression systems, revealing that removing either the extracellular (Δkringle) or the intracellular (Δp85-like) domains abrogated the inhibitory activity of TrIP (16). Furthermore, in the absence of the kringle domain, cells maintained expression of the Flag-tagged intracellular TrIP domain following stimulation, suggesting that the intracellular p85-like domain of TrIP persists for some time. We thus hypothesized that the kringle domain is required for TrIP localization proximal to the TCR, and that cleavage of the extracellular domain enables its exclusion from TCR synapse following activation. More studies are needed to fully define these mechanisms with endogenous TrIP protein, as the currently available tools limit many of these studies to cell line expression systems.

In summary, we present data expanding our understanding of the relationship of PIK3IP1/TrIP to signals that regulate T cell activation. Future studies on the downstream effects of TrIP downregulation in specific disease contexts are still needed to fully understand the impact of TrIP regulation in the T cell compartment. TrIP itself represents a potentially unique biological target, as preventing its inhibition in T cells could improve responses to cancer or infection, whereas strategies to prevent its downregulation may help suppress autoimmunity or chronic inflammation.

## Materials and Methods

### Antibodies and flow cytometry

For extracellular staining, single cell suspensions were stained at 4°C for 30 minutes with Live/Dead + antibody cocktail containing anti-CD16/CD32 (Fc Shield; Tonbo Biosciences: clone 2.4G2) resuspended in PBS. For intracellular staining, cells were fixed/permeabilized using the eBioscience Foxp3/Transcription factor staining buffer set (00-5523-00), as per manufacturer instructions. Following fixation, intracellular staining antibodies were resuspended in 1x permeabilization buffer and used to stain at 4°C for 30 minutes. The following antibodies and dyes were used for these analyses: anti-CD90.2 (BD Biosciences; clone 53-2.1), anti-CD8α (BD Biosciences; clone 53-6.7), anti-CD44 (BD Biosciences; clone IM7), anti-CD11c (BD Biosciences; clone N418), anti-TCRβ (BD Biosciences; clone H57-597), anti-CD4 (BioLegend; clone RM4-5), anti-CD11b (ThermoFisher: clone M170), anti-CD62L (BioLegend; clone MEL-14), anti-F4/80 (BioLegend; clone BM8), anti-CD25 (BD Biosciences: clone 7D4), anti-Gr-1 (Invitrogen: clone RB6-8C5), anti-CD19 (Biolegend: clone 1D3), anti-Foxp3 (Invitrogen; FJK-16s), anti-pS6_(235/236)_ (Cell Signaling; clone D57.2.2E), anti-TCR Vα2 (Biolegend; clone AF-7), anti-CD69 (BioLegend: clone H1.2F3), anti-Flag (BioLegend; clone L5). Live/dead staining was performed using the Zombie NIR fixable viability kit (BioLegend; 423105). For the crispr knockout studies, we additionally used anti-PKCθ (Cell signaling; clone E1I7Y) and anti-ADAM10 (R&D; clone # 139712) for staining. Alexa Fluor 488 Anti-Rabbit IgG was used as a secondary against PKCθ (Jackson ImmunoResearch; Cat No. 711-545-152) and Goat anti-Rat IgG Alexa Fluor 488 (Invitrogen; Cat No. A-11006).

For the majority of TrIP staining experiments, the 18E10 clone showed the most robust and reproducible staining and was therefore used for the studies involving endogenous murine TrIP. We directly conjugated the α-TrIP 18E10 to Alexa Fluor 647 using a commercial labeling kit (Invitrogen A20186), per manufacturer’s instructions. All flow cytometry was performed on a 5-laser Cytek Aurora and flow cytometry data were analyzed with FlowJo (v10.10.0).

### Animals

All experiments involving animals were approved by the University of Pittsburgh Institutional Animal Care and use Committee (IACUC). We previously generated a conditional mouse knockout strain with LoxP sites inserted into the TrIP gene (*Pik3ip1*^*fl/fl*^), targeting the removal of exons 2-5 (16). These mice were crossed to E8i^cre^ mice (Jackson strain #008766), to achieve knockout of *Pik3ip1* specifically in CD8^+^ T cells. For peptide stimulation studies, we employed the P14 TCR transgenic strain that expresses a T cell receptor specific for the gp33 peptide from LCMV (Jackson; Strain #037394). All strains were backcrossed more than 9 generations to C57BL/6J (Jackson; Strain #000664). All mice were bred in-house under SPF conditions, and experimental mice were either littermates or housed in the same facility and same room. All experiments were conducted on mice 6-8 weeks of age, with age and sex-matched groups.

### Generation of TrIP-Ig fusion protein

To generate the TrIP-Ig fusion, the ecto domain of murine TrIP/Pik3ip1 was PCR amplified from IMAGE consortium clone number 4039129, using primers to add EcoRV and BglII sites at the 5’ and 3’ ends, respectively. This fragment was then cloned into the same restriction sites of pFuse-hIgG1-Fc2. Reading frame and integrity of the cloned fragment were confirmed by sequencing. This plasmid was then transfected into HEK293T cells and Protein-A purified by a commercial vendor (SydLabs, Inc.).

### Development of mAb’s to the ecto domain of TrIP

Monoclonal antibodies to the ecto domain of murine TrIP were developed in conjunction with a commercial vendor (Rockland, Inc.). Briefly, Sprague-Dawley rats were immunized with the aforementioned TrIP-Ig fusion protein. Immunized rat splenocytes were then fused with Sp2/0-Ag14 myeloma cells to generate hybridomas. These hybridomas were subsequently subcloned one or more generations via limiting dilution. Reactivity/specificity of each was initially screened via ELISA for reactivity with recombinant TrIP-Ig protein, and counter-screened against human IgG1.

### Generation of TrIP deletion mutant constructs

For the initial antibody validation experiments, full-length human *PIK3IP1* and full-length mouse *Pik3ip1* gBlocks (IDT) were cloned into pcDNA3.1 via restriction enzyme cloning. For TrIP cleavage experiments, two truncated and one full-length TrIP gBlocks (IDT) were produced and cloned into a minimal pcDNA3.1(+) backbone modified from pcDNA3.1(+) (Invitrogen). In the first TrIP truncation, a 16 amino acid sequence was removed [AA positions 133-148; (DNA sequence: CAGTCAGCTTGTGAGGATGAACTCCAAGGAAAAAAAAGACCTAGGAAC)]. For the second truncation, that deletion was extended an additional 13 amino acids to make a 29 amino acid deletion [AA positions 120-148 removed; (DNA sequence: AGGAGTGAGGCAGCCGAG-GTGCAGCCAGTGATCGGGATCAGTCAGCTTGTGAGGATGAACTCCAAGGAAAAAAAAGACCTAGGAAC)]. Each construct used for transfections was designed to include a Flag tag (DYKDDDDK) for antibody recognition via anti-Flag antibody (clone: L5). All plasmid constructs’ MCS were verified by Sanger sequencing.

### Cell lines, transfections and activation

Human embryonic kidney (HEK) 293T cells were cultured in DMEM containing 10% BGS, 1% penicillin/streptomycin, and 1% L-glutamine. D10.G4.1 mouse T helper cell line (D10 cells) were maintained in 50 U/mL rhIL-2 in complete RPMI (cRPMI) media (cRPMI supplemented with 10% BGS, 1% penicillin/streptomycin, 1% L-glutamine, 50 mM 2-mercaptoethanol, 1x non-essential amino acids, 1% HEPES, and 1% sodium pyruvate). Transfection of HEK293T cells was performed using TransIT-LT1 (Mirus Bio; MIR 2304) per manufacturer instructions. Transfection of D10 cells via electroporation was performed using a Bio-Rad GenePulser Xcell (Square Pulse protocol; 400V, 3 pulses, 0.5 ms pulse length, 0.05 s between pulses). For truncated TrIP expression experiments, pMaxGFP was co-transfected as a positive control (Lonza Biosciences). Activation of D10 cells was via plate-coated anti-CD3 (clone:145-2C11; Tonbo Biosciences 50-201-4837); anti-CD28, (clone 37.51; BioLegend: 102101).

### Mouse tissue processing and in vitro stimulation

Spleens harvested from either WT C57BL/6J or WT P14 TCR Tg mice were mechanically disrupted and filtered to single cell suspensions. Red blood cells were removed via 1x RBC lysis buffer (eBioscience 00-4333-57), according to manufacturer instructions. Splenocyte stimulations were performed in complete RPMI (cRPMI) media (cRPMI supplemented with 10% BGS, 1% penicillin/streptomycin, 1% L-glutamine, 0.05 mM 2-mercaptoethanol, 1x NonEessential Amino Acids, 1% HEPES, and 1% sodium pryuvate). Anti-CD3 (clone:145-2C11; Tonbo Biosciences 50-201-4837); and anti-mCD28, clone 37.51; Biolegend: 102101) were used for plate-coating at the indicated concentrations. For P14 TCR Tg stimulations, three different peptides were selected based on published work by J. M. Boulter *et al*. (23); a high affinity gp33 peptide (sequence: KAVYNFATM), the WT gp33 peptide (sequence: KAVYNFATC), and a peptide with lower affinity, L6F gp33 (sequence: KAVYNLATC).

### Inhibitors

For the *in vitro* TrIP downregulation experiments, inhibitors used were as follows: IC87114 (SellickChem; S1268), 10 μM; MK-2206 (Cayman Chem; 11593), 2.5 μM; rapamycin (SellickChem; S1039), 100 nM; dasatinib (Stemcell; 73082), 2.5 nM; FK-506 (SellickChem; S5003), 10 μM. U0126 (MedChemExpress; HY-12031), 2.5 μM; bisindolylmaleimide I (SellickChem; S7208), 10 μM; sotrastaurin (SellickChem; S279101), 5 μM; Go6983 (MedChemExpress; HY-13689), 2.5 μM; Go6976 (MedChemExpress; HY-10183) 1.25 μM GW280264X (Bio-Techne; Cat# 7030), 1 μM; GI254023X (Selleckchem; S8660), 25 μM, TAPI-I (Selleckchem; S7434) 25μM. Cells were pre-treated with inhibitor for 30 minutes prior to peptide stimulation, with each stimulation occurring as a reverse timecourse, to ensure that each sample was subjected to exposure to the drugs for the entirety of the stimulation, totaling 4.5 hrs of drug exposure for all no-stimulation (0hr) and stimulated conditions (4hr).

### CRISPR/Cas9 gene knockout in naïve CD8 T cells

WT CD8^+^ T cells were bead-isolated from splenocytes processed to single-cell suspensions as described above. CRISPR protocols were adapted from work previously described (31, 32). We used magnetic bead based negative selection to enrich for CD8^+^ T cells (>90%), which were at 2×10^6^/mL in complete RPMI (cRPMI) supplemented with 5 ng/mL recombinant IL-7 (PeproTech: Cat. No. 217-17) for 24 hours at 37° C prior to RNP nucleofection, to achieve knockout in naïve cells without prior stimulation. All CRISPR reagents were sourced from IDT. The crRNAs used are as follows; Mm.Cas9.ADAM10.1.AC - Alt-R™ CRISPR-Cas9 crRNA, 10 nmol – (/AltR1/rArG rUrUrC rArArC rCrUrA rCrGrA rArUrG rArArG rGrUrU rUrUrA rGrArG rCrUrA rUrGrC rU/AltR2/), Mm.Cas9.ADAM10.1.AD - Alt-R™ CRISPR-Cas9 crRNA, 10 nmol – (/AltR1/rGrG rUrUrU rCrArU rCrArA rGrArC rUrCrG rUrGrG rGrUrU rUrUrA rGrArG rCrUrA rUrGrC rU/AltR2/), Mm.Cas9.ADAM17.1.AA - Alt-R™ CRISPR-Cas9 crRNA, 10 nmol – (/AltR1/rCrA rUrArU rArCrC rGrGrA rArCrA rCrGrU rGrUrU rUrUrA rGrArG rCrUrA rUrGrC rU/AltR2/), Mm.Cas9.ADAM17.1.AB - Alt-R™ CRISPR-Cas9 crRNA, 10 nmol – (/AltR1/rCrG rUrCrC rUrGrG rCrArC rCrCrC rGrArC rCrUrC rGrUrU rUrUrA rGrArG rCrUrA rUrGrC rU/AltR2/), Mm.Cas9.PRKCQ.1.AC - Alt-R™ CRISPR-Cas9 crRNA, 10 nmol – (/AltR1/rArU rGrUrC rArCrC rGrUrU rUrCrU rUrCrG rArArU rGrUrU rUrUrA rGrArG rCrUrA rUrGrC rU/AltR2/), Mm.Cas9.PRKCQ.1.AD - Alt-R™ CRISPR-Cas9 crRNA, 10 nmol - (/AltR1/rGrC rGrUrC rArArA rGrGrU rGrCrU rGrUrC rCrCrA rGrUrU rUrUrA rGrArG rCrUrA rUrGrC rU/AltR2/), Alt-R® CRISPR-Cas9 tracrRNA (Cat No. 1072532), Alt-R S.p. Cas9 Nuclease V3 (Cat No. 1081058), Alt-R® CRISPR-Cas9 Negative Control crRNA #1 (Cat No. 1072544), Alt-R® Cas9 Electroporation Enhancer (Cat No. 1075915). RNP formation was performed at a 1:1 equimolar ratio with each tracrRNA independently, to maintain a 1:1.5 Cas9:gRNA molar ratio, which was empirically determined. We thus mixed 100 μM gRNA + 61 μM Cas9 to generate RNP’s, which were pooled immediately prior to electroporation at 1:1 for each gene targeting pair (i.e., PRKCQ.1.AD + PRKCQ.1.AC). Electroporation was performed using Lonza P3 Primary Cell 4D-Nucleofector™ X Kit, using pulse setting “DS 137.” Following electroporation, cells were immediately placed in cRPMI (2×10^6^ cells/mL) supplemented with 5 ng/mL rIL-7 and 20% BGS (Hyclone) to support viability and recovery. T cells were maintained in cRPMI + 5 ng/mL rIL-7 for at least 5 additional days following nucleofection to allow for protein turnover. The absence of prior stimulation necessitated these additional days in culture so that we could assess stimulation-dependent protein degradation, as noted in previous reports (31, 32). Cells were stimulated after day 7 in culture with 3 μg/mL plate coated anti-CD3 (clone:145-2C11; Tonbo Biosciences 50-201-4837) for the time points indicated.

## Data Availability Statement

All data are contained in the manuscript and/or can be shared upon request.

## Acknowledgements

This work was supported by grants from the NIH (F31CA261039 and T32CA082084 to B.M.) and (R01 GM136148 to L.P.K.). We would like to thank Grace Bowman and Edgar Cardona for their help with our mouse genotyping. We would also like to thank the Unified Flow Cytometry Core.

## Author Contributions

B.M., S.R., H.B., and L.P.K. contributed to conceptualization. Funding acquisition contributed by B.M. and L.P.K., Data curation, formal analysis, and validation were performed by B.M., S.R., and U.U. Experiments were performed by B.M., S.R., H.B., L.L., U.U., and A.W. Writing – original draft by B.M., Supervision and manuscript editing were performed by L.P.K.

